# Functional Modeling of Plant Growth Dynamics

**DOI:** 10.1101/190967

**Authors:** Yuhang Xu, Yumou Qiu, James C. Schnable

## Abstract

Recent advances in automated plant phenotyping have enabled the collection time series measurements from the same plants of a wide range of traits over different developmental time scales. The availability of time series phenotypic datasets has increased interest in statistical approaches for comparing patterns of change between different plant genotypes and different treatment conditions. Two widely used methods of modeling growth over time are point-wise analysis of variance (ANOVA) and parametric sigmoidal curve fitting. Point-wise ANOVA yields discontinuous growth curves, which do not reflect the true dynamics of growth patterns in plants. In contrast, fitting a parametric model to a time series of observations does capture the trend of growth, however these models require assumptions regarding the true pattern of plant growth. Depending on the species, treatment regime, and subset of the plant lifecycle sampled this assumptions will not always hold true. Here we introduce a different approach – functional ANOVA – which yields continuous growth curves without requiring assumptions regarding patterns of plant growth. We compare and validate this approach using data from an experiment measuring growth of two maize (*Zea mays* ssp. *mays*) genotypes under two water availability treatments over a 21-day period. Functional ANOVA enables a nonparametric estimation of the dynamics of changes in plant traits over time without assumptions regarding curve shape. In addition to estimating smooth curves of trait values over time, functional ANOVA also estimates the the derivatives of these curves – e.g. growth rates – simultaneously. Using two different subsampling strategies, we demonstrate that this functional ANOVA method enables the comparison of growth curves between plants phenotyped on non-overlapping days with little reduction in estimation accuracy. This means functional ANOVA based approaches can allow larger numbers of samples and biological replicates to be scored in a single experiment given fixed amounts of phenotyping infrastructure and personnel.

## Introduction

One of the primary goals of both classical and quantitative genetic research is to link genotypic variation to phenotypic variation by identifying specific genetic variants that produce defined changes in phenotype. In the last several decades, advances in DNA sequencing have drastically increased the throughput and decreased the cost of quantifying genotypic variation across individuals. Today the vast majority of the time and cost of plant genetic research is devoted to capturing and quantifying phenotypic data, a process that remains slow and both cost and labor intensive. The bottleneck of phenotypic data collection has driven interest in automated and high throughput approaches to collecting plant phenotypes. High throughput plant phenotyping platforms use cameras or other sensors to capture non-destructive measurements of plant traits from dozens to thousands of plants per day^1^. Because these measurements are both automated and nondestructive, the same traits can be measured from the same plants repeatedly throughput the life cycle of a plant. Unlike single time point measurements, time series trait data enables the quantification of the dynamics of plant growth and development. Biomass data collected from maize RIL and association populations have demonstrated that different genetic loci are identified using data from different time points in development^2, 3^. However statistical approaches for both dealing with the particular complexities of time series phenotypic measurements and extracting as much information as possible from repeated phenotypic measurements remains an ongoing area of development within plant biology and quantitative genetics.

One approach to dealing with high density time series data is to conduct independent quantitative trait loci (QTL) or association analyses at each individual time point measured^4, 5^. Under the ANOVA setup, we name this method point-wise ANOVA, which performs ANOVA at each time point individually. However, this approach generally requires that all plants be scored at all time points analyzed. In addition it does not leverage the potential of repeated measurements to increase the accuracy with which true values at a given time point can be estimated. Another approach is to fit particular functions such as logistic curves to the data^6, 7^. However, this parametric inference approach will produce accurate results only if the assumptions of the growth model function are satisfied by the observed data. Commonly used growth curve models (sigmoidal curves) generally require data from across the entire life cycle of the plant, which can limit the types of phenotyping data to which these models can be applied. For example, many greenhouse or ground-based phenotyping systems can only be employed to gather data from plants below a fixed height limit^8,9^. For taller crops such as maize or bioenergy sorghum, only a portion of the lifecycle can be phenotyped without exceeding these height limits.

Functional data analysis (FDA)^10,11^, is another approach which can be applied to the analysis of time series phenotypic datasets. This alternative approach combines many of the strengths of both point-wise ANOVA and parametric modeling approaches to the analysis of time series phenotypic datasets. In FDA, data-driven nonparametric approaches^11‒14^ are used to fit the trend of a data series over time. Unlike point-wise ANOVA, FDA makes very flexible assumptions about the distribution of time points^10^. Multiple observations taken from the same plant over time will show a degree of correlation, and if correctly harnessed these correlations can be used to increase the accuracy with which different effects can be estimated. However this correlation structure is often missed or captured incorrectly by time series analysis. In FDA, a mixed random effect term^10^ is used to explain the correlation structure among the data. Statistical inference can also be used to obtain confidence bands for the estimated curves, again taking into account the temporal dependence of the data. FDA has been applied to the analysis of plant phenotypic data in several recent cases. For example, FDA has been used to analyze different levels of variation in root gravitropism data^15^; dominant variation in phenotype data has been extracted by FDA and applied to further analysis, such as multivariate QTL mapping^16^.

In studies aimed at comparing genotypes or treatments, optimal experimental design emphasizes collecting measurements as close to simultaneously as possible for all plants within the study to avoid increased variance across measurements resulting from both developmental and diurnal changes in the measured phenotype. However, in larger quantitative genetic studies using high throughput phenotyping technologies this requirement for simultaneous data collection can become a major bottleneck limiting the number of plants and number of accessions which can be included within a single experiment. For example, the UNL Greenhouse Innovation Center has the capacity to image approximate 400 plants per day, while significantly more total plants can be grown in parallel^17^. Similar systems such as the Bellweather phenotyping system also have the capacity to grow more plants simultaneously than can be imaged over the course of a single day^9^. Phenotypes collected from unmanned aerial vehicles (UAVs) suffer from a similar constraint on how many plots can be imaged per day, with the additional constraint that unsuitable weather conditions – high wind, thunderstorms etc – can result in missing data from particular sites on particular dates, producing unbalanced final phenotypic datasets. In many cases, FDA can provide a way to address this issue by permitting the reconstruction of growth curves using relatively small numbers of measurements spaced across a large period of development, thus generating predicted values for any time points not scored. Here we show that our proposed method can produce a sub-sample estimator, based on half of the observed data for individual plants with only minimal decreases in accuracy relative to estimates constructed from the entire dataset. In addition, we demonstrate accurate estimator values when different batches of plants are phenotyped on alternating days relative to each other.

## Results

The plant high throughput phenotyping datasets used in this study were taken from a factorial experiment with 60 plants divided equally into two genotypes (B73 and FFMM-A) and equally into two treatments (well watered and drought stressed)^17^. The two genotypes were selected because B73 is a widely used reference genotype which is a typical representative of moderate temperate maize, while FFMM-A represents one extreme end of the distribution of plant life cycle speed and plant architecture present within domesticated maize. Plants were imaged daily for a total of 21 days, excluding day 16 as a result of a technical failure. As previously reported, B73 grew faster and larger than FFMM-A, and well watered maize grew faster and larger than drought stressed plants (Figure 1)^17^.

**Figure 1.**
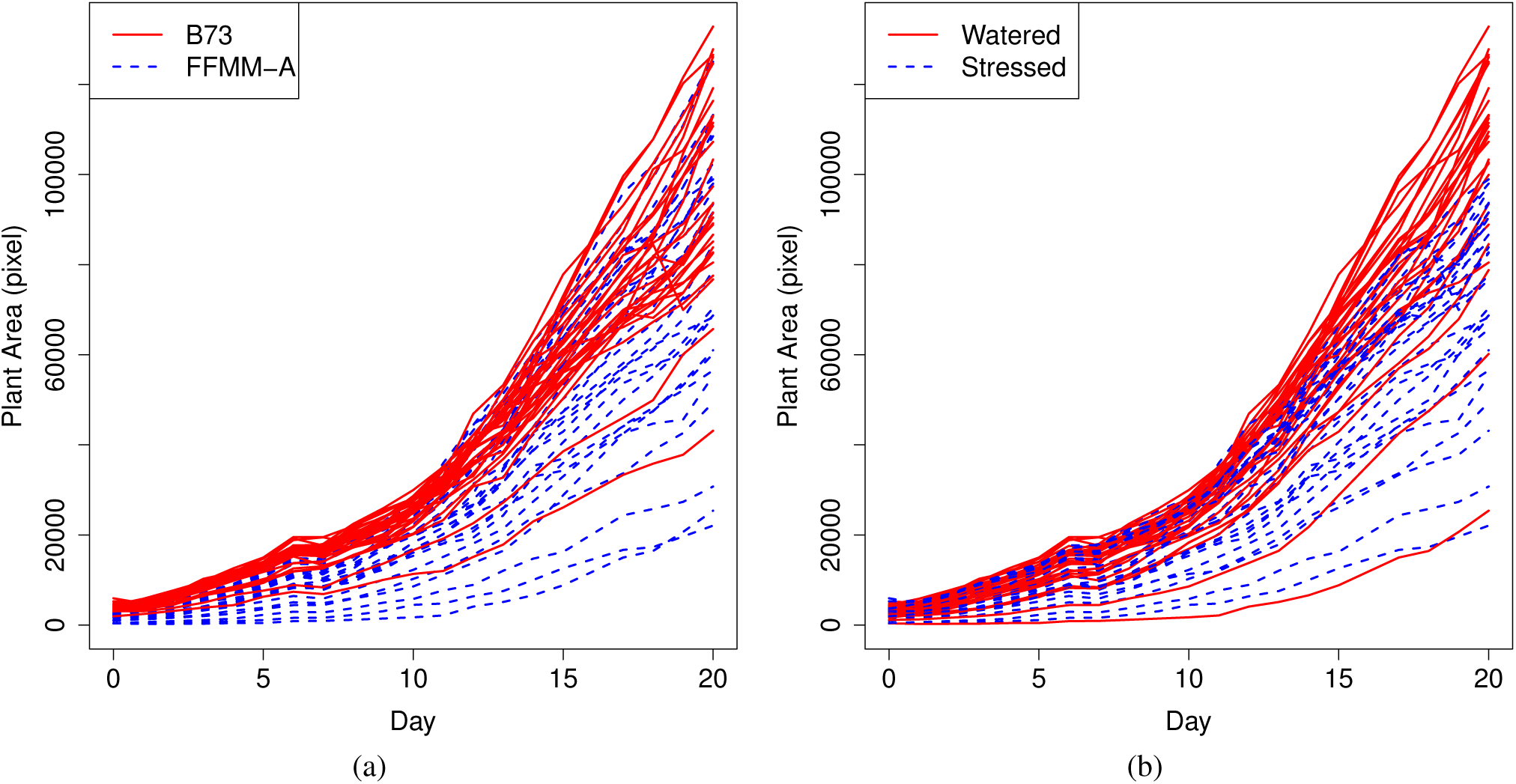
Estimated plant size for each individual phenotyped within this dataset. (A) Plants classified based on genotype. (B) Plants classified based on water treatment. Day 0 corresponds to six days after planting. Day 20 corresponds to day 26 after planting.

### Estimating genotype and treatment effects using spline fitting

Vegetative biomass accumulation in maize and many other crops is generally assumed to follow a sigmoidal growth curve^18^. The cumulative increase of total carbon fixed as the plant produces additional leaves enabled the growth of either more or larger leaves creating the acceleration portion of the growth curve, while later in development much carbon is devoted to reproductive development, slowing the accumulation of additional vegetative biomass which ultimately plateaus, producing a final S-shaped curve. The dataset employed here did not extend into reproductive development and thus captured only the first phase of the sigmoidal biomass accumulation pattern, producing J-shaped curves as shown in Figure 1.

Applying penalized spline smoothing to the data, we obtained the estimated growth under different conditions shown in Panel (a) of Figure 2. As expected, for each genotype, well watered plants are consistently larger than drought stressed plants. In addition, plants from the accession B73 were consistently larger than those of FFMM-A. At early stages in plant development, genotype played a larger role in determining plant biomass than did water treatment. From day 1 to day 16, both well watered and drought stressed B73 were consistently larger than FFMM-A plants in either water treatment. After day 16, the biomass of well watered FFMM-A exceeds that of drought stressed B73.

**Figure 2.**
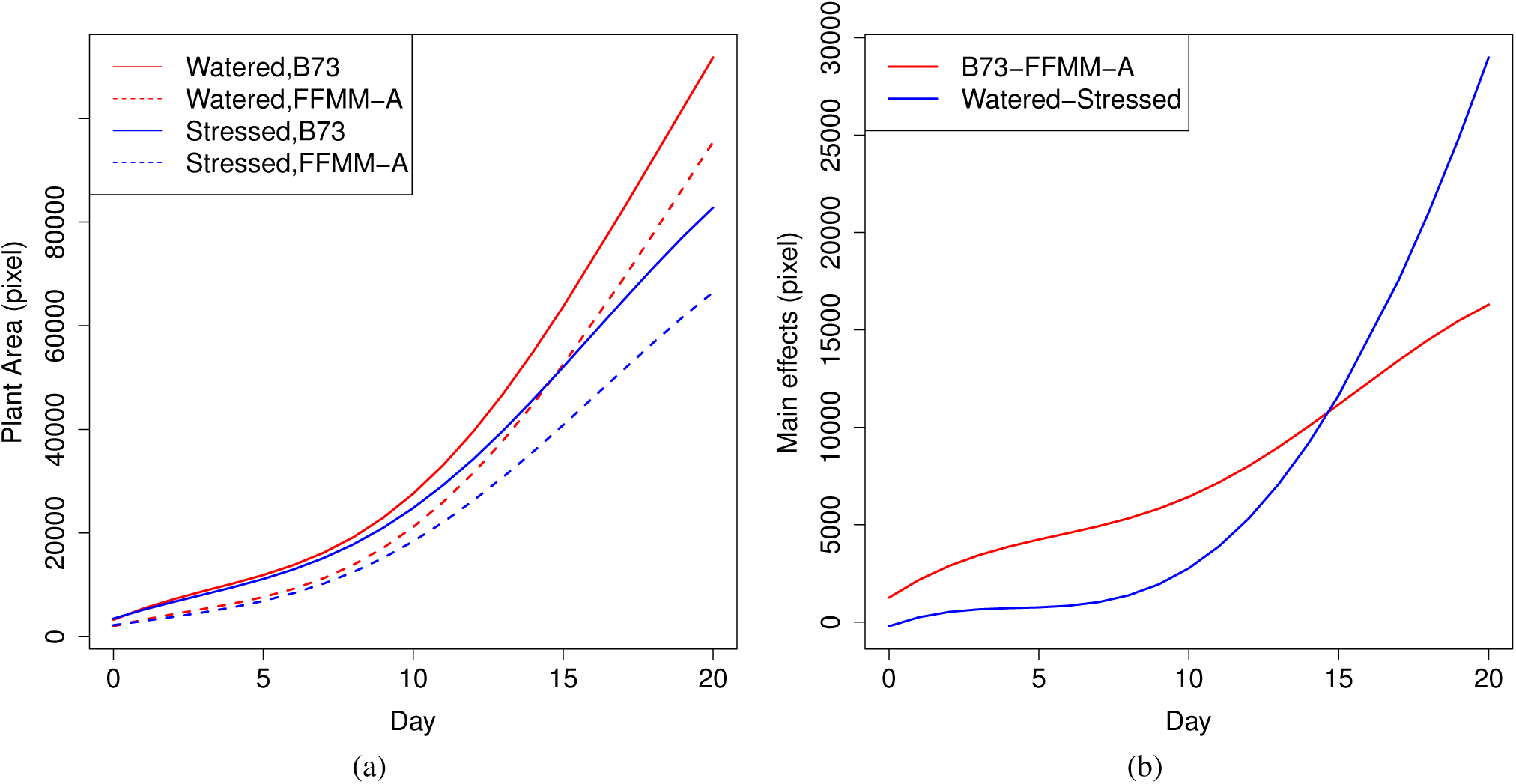
(A) Growth curves estimated for each genotype x treatment combination. (B) Estimated effect sizes for genotype (red) and treatment (blue).

Panel (b) of Figure 2 shows the estimated main effect functions for both genotype and water treatment. Both effect functions are monotone increasing, but they exhibit very different shapes. The effect function for genotype is close to linear and increases steadily. However the effect function for water treatment shows an obvious “J” shape, starting at a low value and increasing very slowly for the first third of the experiment and then growing rapidly. The estimated treatment effect function is close to zero during the first few days because the drought stress started from day 6. The intersection of the two main effects functions coincides with the finding in Panel (a).

### Dynamic changes in growth rate

The results for the dynamics of the growth curves are summarized in Figure 3. Panel (a) shows the estimated growth velocity functions under different conditions. Similarly, for each genotype, the growth velocity for non-water stressed maize is consistently higher than the drought stressed maize; within each water treatment, B73 consistently grows faster than FFMM-A. All growth velocity curves show an “S” shape: during the early period, the growth rates decrease slightly; during the middle period, the growth rates increase sharply; for the later period, the growth rates decline again. Interestingly, the early period for B73 is about two or three days longer than FFMM-A and the late period of non-water stressed maize is about two or three days longer than drought stressed maize. From the velocity perspective, this coincides again with the finding that the genotype effect plays an important role during the early period but the treatment effect plays an important role during the late period. Panel (b) shows the estimated main effect velocity functions. Similarly, the two main effect functions show different shapes: after the early period of decrease, the genotype effect on the rate of growth increases slightly followed by an decrease, but the watering effect on the rate of growth increases sharply and keeps increasing.

**Figure 3.**
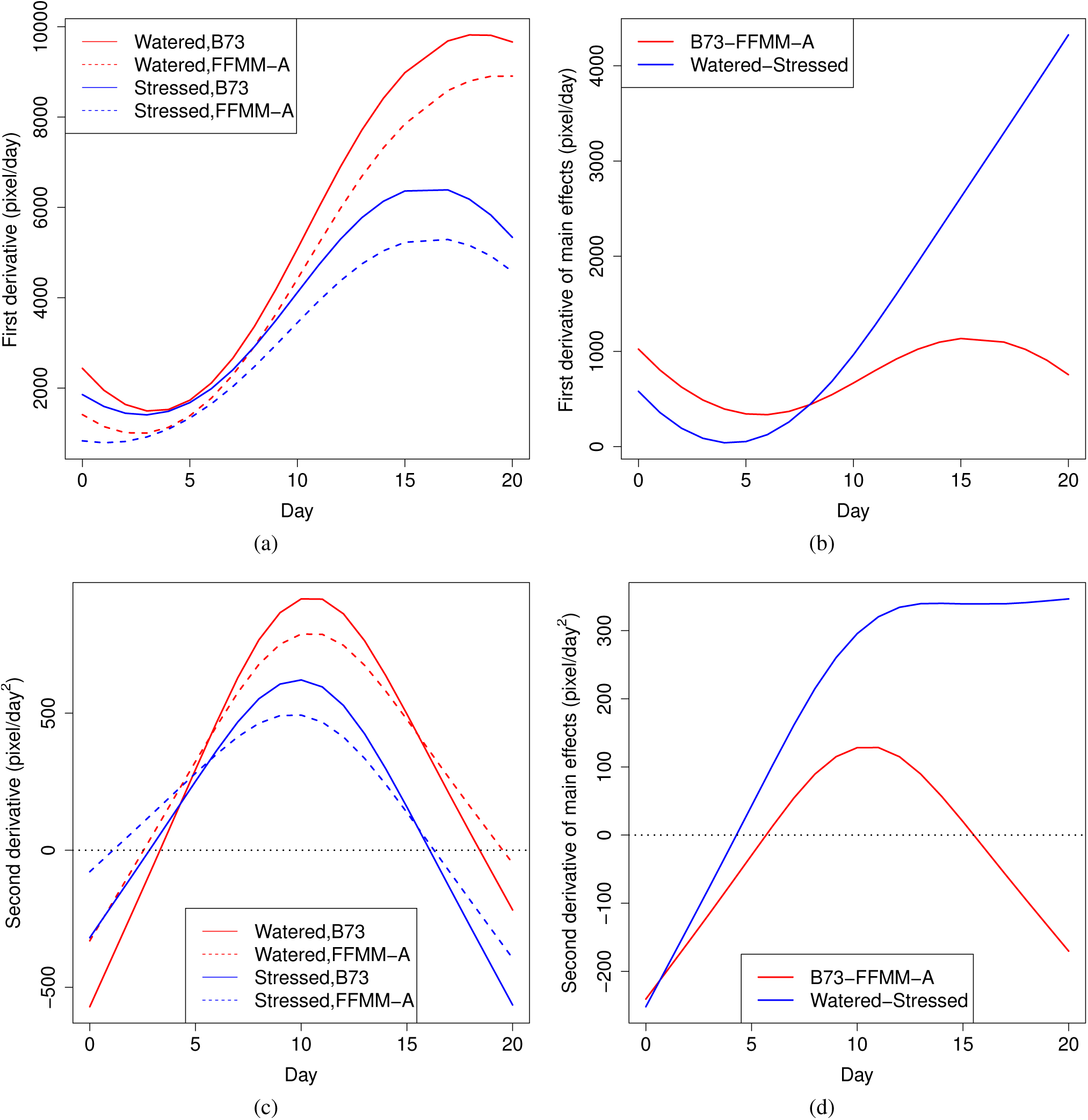
Panel (a) shows the estimated first derivative of growth curves; panel (b) shows the estimated first derivative of main effect function of genotype and water treatment; Panel (c) shows the estimated second derivative of growth curves; panel (d) shows the estimated second derivative of main effect function of genotype and water treatment.

Panel (c) shows the estimated growth acceleration functions under different conditions. Each curves in panel (c) exhibits a parabola-like shape with the maximum acceleration located around the tenth day of the experiment. Panel (d) shows the estimated main effect acceleration functions. The treatment effect of the acceleration of growth seems consistently higher than the genotype effect except the first few days. Both acceleration functions increase during the first half of the experiment. However, for the later half period, the watering effect on acceleration becomes close to a constant, about 340, whereas the genotype effect on acceleration decreases dramatically.

### Functional ANOVA for comparing growth with non-overlapping time points

To investigate the estimation efficiency when plants were not phenotyped every day and to test the prediction accuracy of functional ANOVAs for plant areas when phenotype measurements were not recorded, comparisons via cross validation were made between the full dataset and subsampled datasets.

The data were subsampled in two ways. In the first scenario, only measurements from odd numbered days were retained (10 days in total) for all the plants. This subsampling tests the effect of reducing the number of days of imaging for a single experiment, allowing more independent experiments to be conducted in parallel using the same infrastructural capacity for phenotypic data acquisition. In the second scenario, all the plants were equally divided into two groups among the two genotypes and two treatments. For the first group, only measurements from odd-numbered days were retained, while for the remaining plants in the second group, only measurements from even-numbered days were retained. This subsampling tests the effect of measuring different subsets of plants in an experiment at different time points, which allows experiments with large number of genotypes or large sample sizes within each genotype to be conducted given a fixed facility capacity for phenotypic data acquisition.

Growth rates together with genotype and treatment effects were estimated using the two subsampling approaches described above by the same functional ANOVA procedure for the full dataset. With the exception of the first several days when all the plants were quite small, as shown in Figure 4, functional ANOVA with half of the data produced reliable estimates, which were within 5% deviation from the estimation using the entire data set. Comparing to the results when all the plants were phenotyped everyday, the average relative estimation difference caused by reducing the imaging frequency was 1.64% (scenario # 1), and that caused by decreasing the number of daily phenotyped plants was 1.45% (scenario # 2). Given those small estimation differences, the functional ANOVA approach is able to recover the entire genotype and treatment effects over time even if the plant iamges are recorded only on half time of the whole experiment or only half of those plants are imaged everyday. This advantage of functional ANOVA provides a guideline on designing larger experiment with more genotypes and parallelizing several experiments to run simultaneously, which could improve the efficiency of the high throughput phenotyping facility.

**Figure 4.**
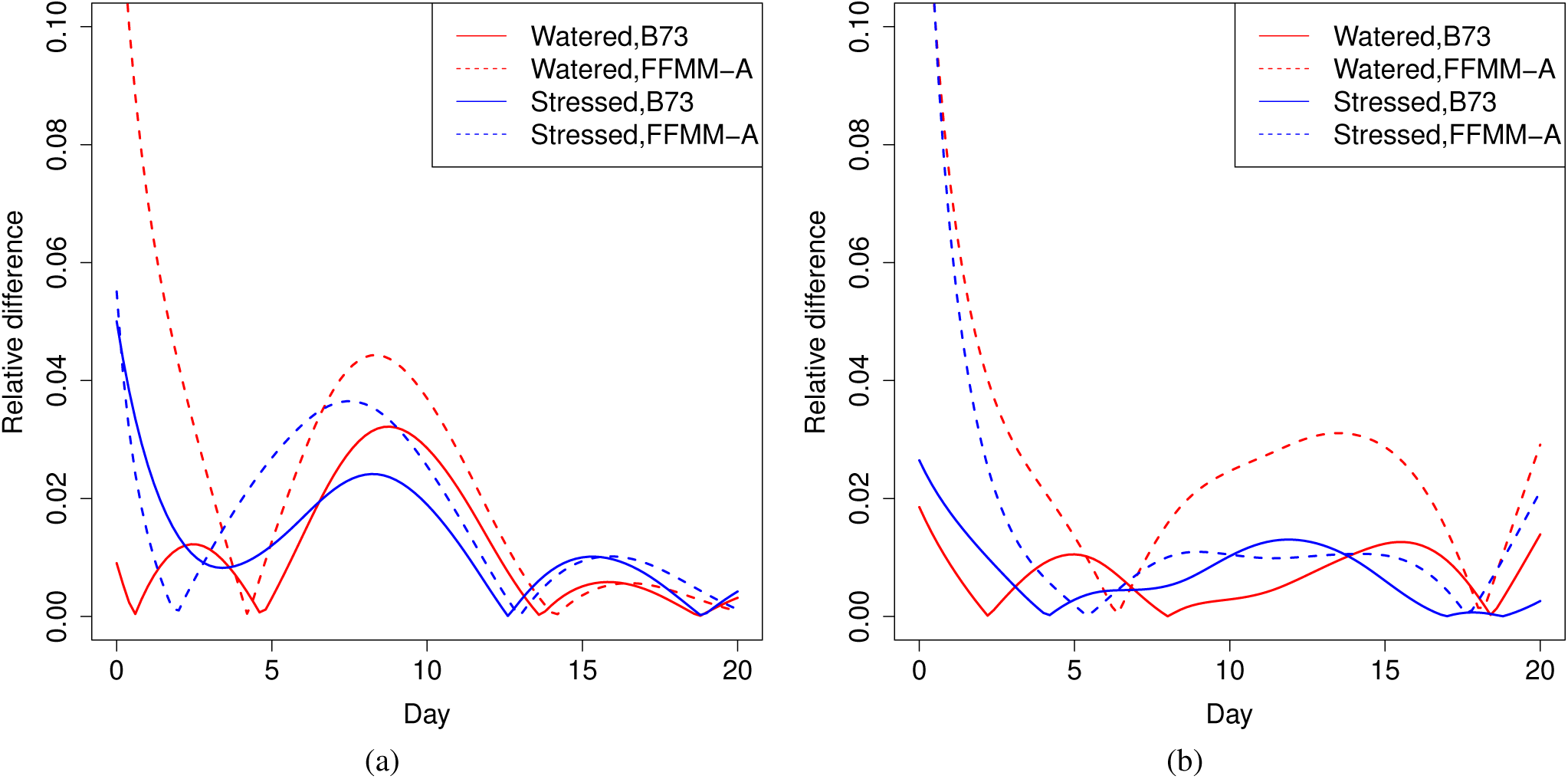
Panel (a) shows the relative difference in estimated growth curves between the whole dataset and a dataset subsampled to include data from only every second day (all plants measured on the same days); panel (b) shows the relative difference in estimated growth curves between the whole dataset and a dataset subsampled to include data from only every second day (plants split into two groups measured on alternating days).

## Discussion

Here we used functional data analysis as a nonparametric method to model plant growth over time. This nonparametric approach effectively incorporates neighborhood information when fitting the underlying growth curve, and produces more accurate estimates of genotype and treatment effects over time^11^. Intuitively, plant biomass at time *t*_0_ is highly related to that at the previous time point *t*_0_ – 1 and the following one at *t*_0_ + 1. This sharing of data between nearby time points also provides increased accuracy for predictions of plant traits at time points not sampled in the experiment. Unlike parametric approaches, the functional data analysis method outlined above is data-driven rather than model driven and thus applicable to a wider range of treatments, genotypes, and developmental stages and adaptive to temporal dependent observations. Compared to parametric modeling approaches, nonparametric methods such as the one employed here are flexible with regards to patterns of grown which do not match prior assumptions regarding the growth pattern of plants. In addition they adjust for the temporal dependence effect in statistical inference, which is generally not considered in parametric approaches.

Finally, we demonstrate that our proposed method is robust to missing data and nonoverlapping sampling dates between subsets of samples within a single experiment. The necessity to collect measurements from all or nearly all individuals at each time point within an experiment is a major constraint on high throughput phenotyping studies in both the greenhouse – where plant measurements are limited by the throughput of imaging systems – and field – where plant measurements are limited by the availability of human labor and weather suitable for phenotyping. The wider adoption of functional data analysis in the analysis of plant phenotyping data, and the awareness of the increased flexibility it provides for sampling datas within experimental designs should lead to larger and most statistically robust experiments in the future.

## Methods

### Experimental Design, Growth Conditions, And Imaging

B73 plants were grown from a seed source validated using RNA-seq SNP calling to match the B73 genotyped used to generate the maize reference genome^19^. Fast Flowering Mini-Maize-A seeds were provided by Morgan E. McCaw and has also been subjected to 24x whole genome resequencing^20^. All plants were grown at the UNL Greenhouse Innovation Center. Plants were sown into 5.7L pots with Fafard germination mix and watered to a target weight of 5.4 kilograms. From DAP 6 (days after planting) to DAP 26, plants were imaged using an RGB camera from angles offset from each other by 90 degrees. Until DAP 10, each plant was rewatered to a target weight of 5.4 kilograms. From DAP 11 (the 6th day since the beginning of imaging) to the end of the experiment drought treated plants received no additional water, while well watered plants continued to be rewatered to a target weight of 5.4 kilograms each day. Further details on experimental design and growth conditions is provided in the reference^17^.

### Extraction of Pixel Counts From RGB Images

A RGB image processing procedure^17^ is applied to extract plant sizes from the acquired images. A threshold is applied on the contrast of green intensity and the average intensity of red and blue to seperate the plant pixels from the background. The majority of the background in our imaging chamber is white. Therefore, the plant areas can be obtained efficiently by such a comparison. The total pixel counts of the extracted plant are considered as a measurement of the plant size.

### Point-wise ANOVA Model

Let *y*_*i*_(*t*_*j*_) be the area of the ith maize measured at time *t*_*j*_, where *i* = 1,…, *n*, *n* = 60 is the sample size, and *j* = 1,…, *m*, *m* = 20 is the number of measured days. Define genotype indicator *G*_*i*_ as follows: *G*_*i*_ = 1 if the ith maize is of genotype B73 and *G*_*i*_ = 0 if the ith maize is of genotype FFMM-A. Similarly, define the environment indicator *W*_*i*_ as follows: *W*_*i*_ = 1 if the ith maize is well watered and *W*_*i*_ = 0 if the ith maize is water stressed. A natural way to model the growth over time is to use the following point-wise ANOVA model:

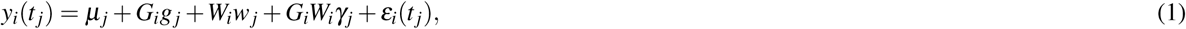

where *μ*_*j*_ is the plant area of water stressed FFMM-A maize at time *t*_*j*_, *g*_*j*_ is the genotye effect function at time *t*_*j*_, *w*(*t*) is the treatment effect function at time *t*_*j*_, *γ*_*j*_ is the genotype–environment interaction at time *t*_*j*_, and *ε*_*i*_(*t*_*j*_) is a zero-mean random variable.

It is interesting to know whether the genotype–environment interactions exist. To explore this, we tested the genotype–environment interaction point-wisely and summarized the results in Table 1. Since all genotype–environment interactions manifested to be insignificant, we revised (1) and used the following point-wise ANOVA model:

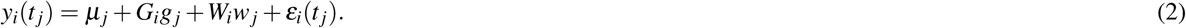

Denote the resulting estimates as 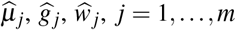. Interpolating the corresponding estimates yields Figure 5. Note that for point-wise ANOVA, the estimates are fitted at each time point *t*_*j*_, so the obtained growth curves and main effects curves are discontinuous, which does not reflect the continuous growth of plants. Moreover, the dynamics, namely velocity and acceleration, of the growth curves cannot be obtained using the point-wise ANOVA method. For these reasons, in this paper, we advocate the following functional ANOVA method.

**Table 1.**
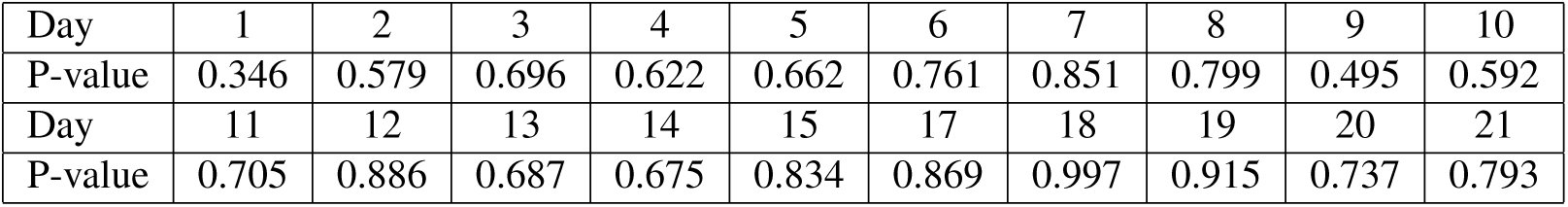
Testing of the genotype-environment interaction point-wisely.

**Figure 5.**
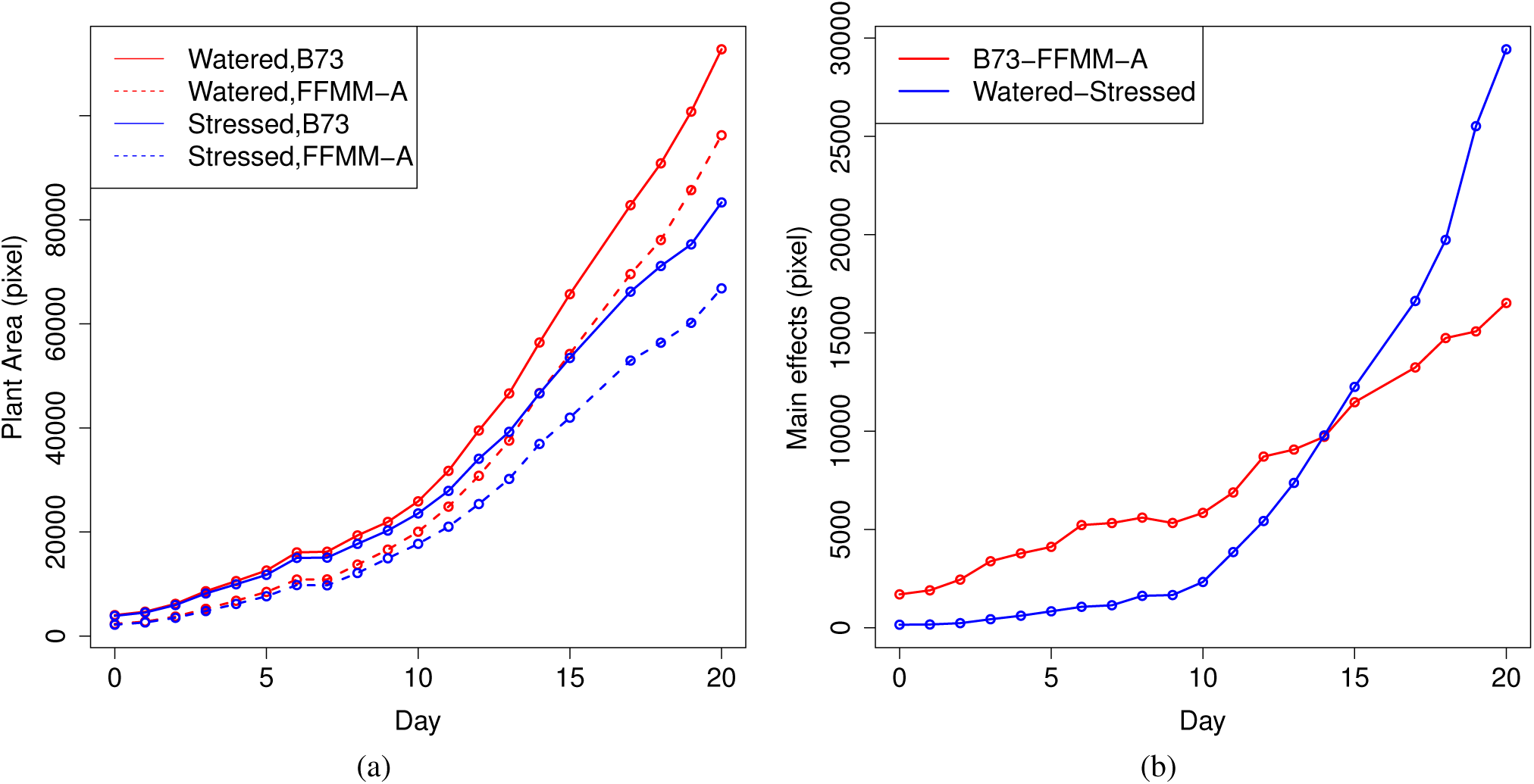
Panel (a) shows the estimated growth curves; panel (b) shows the estimated main effect function of genotype and water treatment by the point-wise ANOVA model.

### Functional ANOVA Model

We assume the following functional ANOVA model for the plant growth

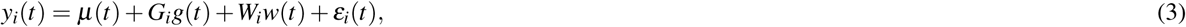

where *μ*(*t*) is the growth function of the water stressed FFMM-A maize, *g*(*t*) is the genotype effect, *w*(*t*) is the treatment effect, and *ε*_*i*_(*t*) is a zero-mean random process. We assume *μ*(*t*), *g*(*t*), and *w*(*t*) are smooth functions. To recover the underlying functions and their dynamics, namely velocity and acceleration, we use penalized smoothing splines^11^.

We first represent *μ*(*t*) using a rank *K* spline basis expansion 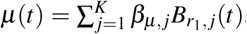 where *β*_*μ*,*j*_ is a coefficient and *B*_*r*_1_,*j*_(*t*) is an order *r*_1_ B-spline basis function. We chose *K* = 12 for a reduced rank representation and let B-spline basis functions have equally-spaced interior knots on [0,20]. Since we are interested in estimating velocity and acceleration functions smoothly, we chose order *r*_1_ = 6. Define ***β***_*μ*_ = (*β*_*μ*, 1_,…, *β*_*μ*, *K*_)^T^ and ***B***(*t*) = (*B*_6,1_,…,*B*_6, *K*_)^T^(*t*). Denote the *r*_2_th derivative of ***B***(*t*) by ***B***^(*r*_2_)^(*t*). Then *μ*(*t*) can be rewritten as *μ*(*t*) = ***B***(*t*)^T^***β***_*μ*_. Similarly we approximate other functions as *g*(*t*) = ***B***(*t*)^T^***β***_*g*_, and *w*(*t*) = ***B***(*t*)^T^***β***_*w*_, To estimate the vectors of parameters, ***β***_*μ*_, ***β***_*g*_, and ***β***_*w*_, penalized smoothing splines minimize the following penalized sum of squares

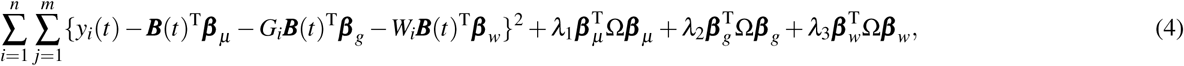

where *λ*_*l*_, *l* = 1,2,3, are smoothing parameters, and **Ω** = ∫***B***^(*r*_2_)^ (*t*){***B***^(*r*_2_)^ (*t*)}^T^d*t* is a penalty matrix. Let *λ* = *λ*_1_ = *λ*_2_ = *λ*_3_ for simplicity, set *r*_2_ = 4 because we penalize the second derivatives. We chose the smoothing parameter *λ* using generalized cross-validation (GCV) and minimized the penalized sum of squares in (4) to obtain the estimates 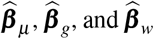. Accordingly, the obtained estimates for the smooth functions are 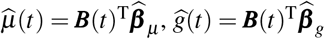, and 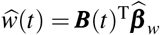. One advantage of using penalized smoothing splines technique is it readily yields different derivatives of the target smooth curves. For example, the estimates of the first and second derivative of *μ*(*t*) are 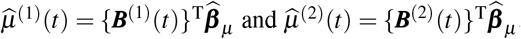, respectively.

## Acknowledgements

The authors wish to acknowledge Yufeng Ge and Piyush Pandey for their willingness to share prepublication data, Yang Zhang for critical evaluation of an early draft of this manuscript and Zhikai Liang for helpful feedback during the initial conception of this experiment.

## Author contributions statement

Y.X., Y.Q. and J.C.S. conceived the experiments. Y.X. and Y.Q. conducted the analysis. Y.X., Y.Q. and J.C.S. wrote the manuscript.

## Additional information

The raw image data employed in this study is hosted at CyVerse under doi 10.7946/P22K7V.x

## References

1. Fahlgren, N., Gehan, M. A. & Baxter, I. Lights, camera, action: high-throughput plant phenotyping is ready for a close-up. Curr. opinion plant biology 24, 93–99 (2015).

2. Muraya, M. M. et al. Genetic variation of growth dynamics in maize (zea mays l.) revealed through automated non-invasive phenotyping. The Plant J. 89, 366–380 (2017).

3. Zhang, X. et al. High-throughput phenotyping and qtl mapping reveals the genetic architecture of maize plant growth. Plant Physiol. 173, 1554–1564 (2017).

4. Moore, C. R. et al. High-throughput computer vision introduces the time axis to a quantitative trait map of a plant growth response. Genet. 195, 1077–1086 (2013).

5. Kwak, I.-Y., Moore, C. R., Spalding, E. P. & Broman, K. W. A simple regression-based method to map quantitative trait loci underlying function-valued phenotypes. Genet. 197, 1409–1416 (2014).

6. Xavier, A., Hall, B., Hearst, A. A., Cherkauer, K. A. & Rainey, K. M. Genetic architecture of phenomic-enabled canopy coverage in glycine max. Genet. 206, 1081–1089 (2017).

7. Deng, J. et al. Models and tests of optimal density and maximal yield for crop plants. Proc. Natl. Acad. Sci. 109, 15823–15828 (2012).

8. White, J. W. & Conley, M. M. A flexible, low-cost cart for proximal sensing. Crop. Sci. 53, 1646–1649 (2013).

9. Fahlgren, N. et al. A versatile phenotyping system and analytics platform reveals diverse temporal responses to water availability in setaria. Mol. plant 8, 1520–1535 (2015).

10. Yao, F., Muller, H.-G. & Wang, J.-L. Functional data analysis for sparse longitudinal data. J. Am. Stat. Assoc. 100, 577–590 (2005).

11. Ramsay, J. O. & Silverman, B. W. Functional Data Analysis, 2nd ed. (Springer-Verlag, New York, 2005).

12. Cleveland, W. S. & Devlin, S. J. Locally weighted regression: an approach to regression analysis by local fitting. J. Am. statistical association 83, 596–610 (1988).

13. Fan, J. & Gijbels, I. Local polynomial modelling and its applications (CRC Press, 1996).

14. Jacoby, W. G. Loess:: a nonparametric, graphical tool for depicting relationships between variables. Elect. Stud. 19, 577–613 (2000).

15. Xu, Y., Li, Y. & Nettleton, D. Nested hierarchical functional data modeling and inference for the analysis of functional plant phenotypes. J. Am. Stat. Assoc. (2017). Http://dx.doi.org/10.1080/01621459.2017.1366907.

16. Kwak, I.-Y., Moore, C. R., Spalding, E. P. & Broman, K. W. Mapping quantitative trait loci underlying function-valued traits using functional principal component analysis and multi-trait mapping. G3: Genes, Genomes, Genet. 6, 79–86 (2016).

17. Ge, Y., Bai, G., Stoerger, V. & Schnable, J. C. Temporal dynamics of maize plant growth, water use, and leaf water content using automated high throughput rgb and hyperspectral imaging. Comput. Electron. Agric. 127, 625–632 (2016).

18. Erickson, R. O. Modeling of plant growth. Ann. Rev. Plant Physiol. 27, 407–434 (1976).

19. Liang, Z. & Schnable, J. C. Rna-seq based analysis of population structure within the maize inbred b73. PloS one 11, e0157942 (2016).

20. McCaw, M. E., Wallace, J. G., Albert, P. S., Buckler, E. S. & Birchler, J. A. Fast-flowering mini-maize: Seed to seed in 60 days. Genet. 204, 35–42 (2016).

